# A biophysical approach to predicting protein-DNA binding energetics

**DOI:** 10.1101/012864

**Authors:** George Locke, Alexandre V. Morozov

## Abstract

Sequence-specific interactions between proteins and DNA play a central role in DNA replication, repair, recombination, and control of gene expression. These interactions can be studied *in vitro* using microfluidics, protein-binding microarrays (PBMs), and other high-throughput techniques. Here we develop a biophysical approach to predicting protein-DNA binding specificities from high-throughput *in vitro* data. Our algorithm, called BindSter, accommodates multiple protein species competing for access to DNA and alternative binding modes of the same protein, while rigorously taking into account all sterically allowed configurations of DNA-bound particles. BindSter can be used with a hierarchy of protein-DNA interaction models of increasing complexity. We observe that the quality of BindSter predictions does not change significantly as some of the energy parameters vary over a sizable range. To take this degeneracy into account, we have developed a graphical representation of parameter uncertainties, called IntervalLogo. We find that our simplest model, in which each nucleotide in the binding site is treated independently, performs better than previous biophysical approaches. The extensions of this model, in which contributions of longer words are also considered, result in further improvements, underscoring the importance of higherorder effects in protein-DNA energetics. In contrast, we find little evidence for multiple binding modes for the transcription factors (TFs) in our dataset. Furthermore, there is limited consistency in predictions for the same TF utilizing microfluidics and PBM experimental platforms.

## Introduction

Sequence-specific interactions between proteins and genomic DNA control numerous cellular processes. An important class of DNA-binding proteins is transcription factors (TFs), which regulate expression of their target genes. Knowledge of *in vivo* binding locations of TFs is important for reconstructing regulatory networks of gene transcription and for understanding which factors control which genes. These locations can be mapped genome-wide using chromatin immunoprecipitation followed by microarray hybridization (ChIP-chip) or sequencing (ChIP-seq) [1, 2]. The latest version of this technique, ChIP-exo, uses exonucleases to trim immunoprecipitated DNA fragments to precise locations of bound proteins, achieving single-base-pair (bp) precision [3]. Many factors influence TF binding locations in living cells, including chromatin structure, which strongly modulates DNA accessibility [4], and cooperative or competing interactions with other DNA-binding proteins [5]. However, the primary determinant of TF binding locations is intrinsic DNA sequence specificity: TFs can recognize and bind their cognate sites on the background of numerous competing sites available in the genome.

Therefore, an accurate *in vitro* picture of intrinsic TF-DNA binding affinity and specificity can provide insights into *in vivo* TF function. For example, high-affinity TF binding sites that are not occupied *in vivo* might indicate that these sites are covered with nucleosomes (the nucleosome is a fundamental unit of chromatin which packages 147 bp of genomic DNA into a tightly bent left-handed superhelix [6]). Conversely, low-affinity sites bound *in vivo* might be a sign of indirect or cooperative binding [7]. Recently, several new technologies have been developed that enable high-throughput *in vitro* determination of protein-DNA binding affinities (see [8] for a recent review). Here we focus on two of these technologies: mechanically induced trapping of molecular interactions (MITOMI) [9, 10], and protein-binding microarrays (PBM) [11, 12].

The MITOMI device uses microfluidics to simultaneously measure binding affinities of a TF to a few thousand DNA probe sequences. The second-generation device [10] contains 4,160 unit cells; each cell has DNA probes with identical sequences, and a given sequence appears in at least two unit cells. All DNA probes are 70 bp long: a probe starts with CGC followed by a 52-bp variable region and a 15-bp fixed sequence at the 3’ end used for labeling and primer extension. Sequences in the variable region are designed to accommodate all 65,536 DNA 8-mers. Two fluorescent labels are used to quantify the number of surface-immobilized protein molecules (BODIPY) and protein-bound DNA probes (Cy5); the ratio of Cy5 to BODIPY fluorescence is linearly proportional to the total protein occupancy of each probe. Fordyce et al. [10] report MITOMI measurements of 28 TFs from *S. cerevisiae* comprising ten different families.

The PBM approach to studying protein-DNA interactions utilizes microarrays with up to tens of thousands of spots. Each spot contains double-stranded DNA probes with identical sequences; the total number of spots is high enough to allow for all possible permutations of a 10 bp-long sequence. DNA probes are fluorescently labeled in order to monitor the consistency of primer-directed DNA synthesis responsible for creating double-stranded DNA at each spot on the array. TF molecules are added to the array and the array is subsequently washed to remove weakly bound and unbound molecules. Finally, the number of remaining bound proteins is quantified using a fluorescent antibody (Alexa488). The antibody fluorescence intensity (normalized by the DNA fluorescence intensity at each spot; DNA is labeled with Cy3) is then proportional to the total protein occupancy of a given probe sequence.

Here we present an algorithm, called BindSter, for inferring energetics of protein-DNA interactions from high-throughput measurements such as MITOMI and PBM, which report total protein occupancies for a set of DNA probe sequences. Under the assumption of thermodynamic equilibrium, these occupancies can be computed using efficient recursive techniques [13, 14] for an arbitrary protein-DNA interaction energy model. The parameters of the model that determine binding energy can then be fit against the data. We consider all sterically allowed configurations of proteins bound to each probe; multiple binding events, overlapping sites, and multiple DNA-bound species (including alternative binding modes of the same TF) are treated rigorously and do not entail significant computational costs. Thus our framework is more consistent in treating steric exclusion and multiple-species competition for DNA sequence compared to previous biophysical models of protein-DNA energetics: MatrixREDUCE [15] and BEEML-PBM [16, 17].

Moreover, our approach is designed to test a hierarchy of protein-DNA energy models of increasing complexity. We start with a basic mononucleotide model in which the contribution of each nucleotide in the binding site to the total interaction energy is independent of all the other nucleotides. This assumption is commonly used in constructing position-specific weight matrices (PWMs) from TF binding site data [18]. Next, we extend the mononucleotide model in two distinct ways by including: (a) dinucleotide contributions; (b) energies of di-and trinucleotides regardless of their position within the binding site. In addition, we check for alternative TF binding modes by fitting two models to the data either simultaneously or sequentially. Although we have focused on 28 TFs with MITOMI data in this work, we have also made predictions for 12 TFs using PBM data, and have carried out a detailed comparison of PBM- and MITOMI-derived models. The software for our algorithm is available online at: http://nucleosome.rutgers.edu/nucenergen/bindster/.

## Materials and Methods

### Physical model of protein-DNA interactions

Consider an ensemble of *M* species of particles distributed along a DNA segment of *N* bp in length. The particles can bind anywhere within the DNA segment subject to steric exclusion (adjacent particles cannot overlap). Partially bounds states (e.g. where the protein hangs off the DNA) are also not allowed. Proteins may bind to either DNA strand. A grand-canonical partition function for this system of DNA-bound particles is given by:

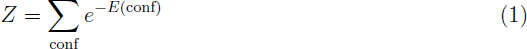

where “conf” denotes an arbitrary configuration of DNA-bound, non-overlapping particles, and *E*(conf) is the total DNA-binding energy (for simplicity we set *k*_*B*_*T* = 1, where *k*_*B*_ is the Boltzmann constant and *T* is the temperature; note that chemical potentials are included implicitly into *E*(conf)).

One can compute *Z* efficiently using a recursive relation:

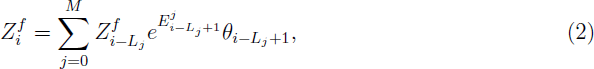

with the initial condition 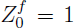 (*i* = 1 … *N*). Here, 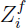 is the forward partial partition function, 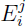 is the binding energy for a species *j* particle at position *i*, *L*_*j*_ is the length of the DNA footprint for a species *j* particle, and *θ*_*i*_ is the Heavyside step function which is equal to 1 when *i* is positive and 0 otherwise. The zeroth species represents the background (no particle), with *L*_0_ = 1 and 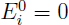 for all *i*.

Likewise, we can compute the partial partition functions in the reverse direction:

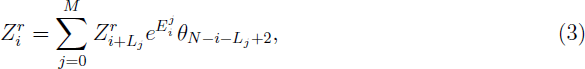

with the initial condition 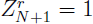 (*i* = *N* … 1). Note that 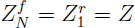 by construction. Further, the probability of starting a particle of species *j* at position *i* is given by:

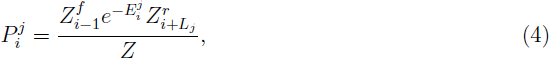

where *i* = 1 … *N* − *L*_*j*_ + 1. Note that 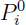 gives the probability of finding no particle at position *i*. The expected number of particles of species *j* covering bp *i* (particle occupancy 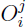) is then given by 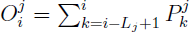 (*i* = 1 … *N*). Finally, the average total number of particles on the entire DNA segment 〈*N*〉 is given by

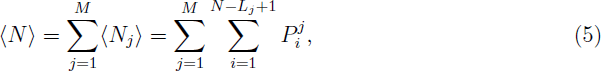

where 〈*N*_*j*_〉 is the average total number of particles of species *j*. Correspondingly, 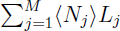 yields the total occupancy of the DNA segment. Taking binding to both strands into account amounts to replacing energies 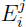 with “free energies” 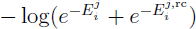, where 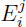 is the binding energy of a particle of species *j* to the *i* … *i* + *L*_*j*_ − 1 site on the upper strand, and 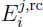 is the binding energy of a particle of species *j* to the *i* + *L*_*j*_ − 1 … *i* site on the lower strand.

We have also tested a simplified DNA-binding model which lacks steric exclusion. In this model, binding of each particle at a given position is independent of all the other particles. The probability of binding 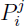 is then simply

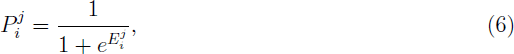

and the quantities 〈*N*〉 and 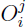 are calculated as above.

### Models of protein-DNA binding energy

#### Mononucleotide model

The mononucleotide model assumes independent contributions to the total protein-DNA interaction energy from each nucleotide in the binding site:

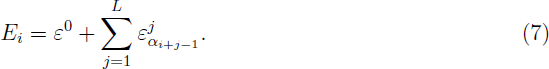

Here, *E*_*i*_ is the total energy of binding at position *i* on a DNA segment (we suppress the species label for brevity), *L* is the binding site length, *α*_*k*_ = {*A*, *C*, *T*, *G*} is the nucleotide at position *k* within the DNA segment, 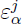 is the energy contribution from the base *α* at position *j*, and *ε*^0^ is a sequence-independent offset which subsumes the chemical potential. We constrain the 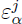 parameters at each position *j* so that 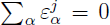, ensuring that the set of fitted parameters is non-degenerate [14]. With these constraints, the mononucleotide model has 3*L* + 1 independent parameters.

#### Dinucleotide model

The dinucleotide model includes both mono- and dinucleotide energy contributions:

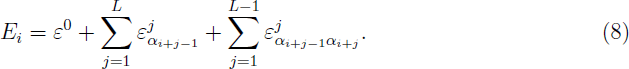

Here, 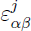 represents the dinucleotide contribution from dinucleotide *αβ* at position *j*, *j* + 1; all other symbols are as in Eq. 7. The mononucleotide parameters are constrained as before, and the dinucleotide parameters are constrained so that 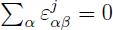 and 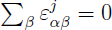 for all *j*. Thus the dinucleotide model has 3^2^(*L* − 1) + 3*L* + 1 = 12*L* − 8 independent parameters. *k*-**mer model**. Another extension of the mononucleotide model adds energy contributions of longer DNA words, irrespective of where these words occur within the binding site. In this model, the binding energy is given by

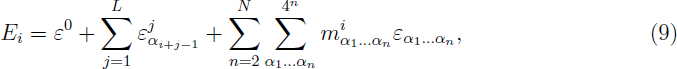

where *N* is the maximum length of DNA words (2 or 3 in our fits), *α*_1_ … *α*_*n*_ is a DNA word of length *n*, 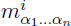 is the number of times that word appears within the binding site, and 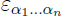 is the energy contribution associated with that word. The energies are constrained by

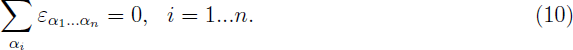

With *N* = 3, there are 3^2^ + 3^3^ = 36 independent *k*-mer parameters in addition to 3*L* + 1 mononucleotide parameters.

#### Position-independent (PI) model

We have also constructed a model where contribution of a given DNA word is independent of its position within the binding footprint:

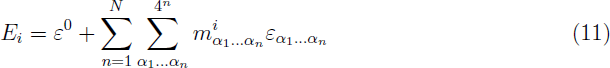

All symbols are as in Eq. 9, and the energy parameters are constrained by Eq. 10. We use *N* = 2 and *N* = 3 with this model.

#### Alternative motif models

These models are designed to account for protein binding in two distinct sequence-dependent modes. The binding energy for one of the modes is given by the mononucleotide model (Eq. 7). In the two-motif model, the other mode is also described by the mononucleotide model, and both models are fit simultaneously. In the secondary motif model, one of the mononucleotide motifs is fit first and then held fixed (except for the *ε*^0^ term), while the parameters of the “secondary” mononucleotide model are allowed to vary. Finally, in the “mononucleotide + PI” model the alternative binding mode is described by the PI model (Eq. 11), which is fit simultaneously with the mononucleotide model.

### PBM location bias

A known source of bias in PBM experiments is the increasing difficulty for the proteins to bind at positions closer to the glass slide [19]. We account for this effect by introducing a bias “energy”, modeled as a quadratic function of the absolute position on the DNA segment:

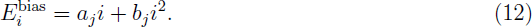

Here, *i* is the position along the DNA probe, and *a*_*j*_ and *b*_*j*_ are fitting parameters. The bias energy can be added to any of the protein-DNA energy models described above, separately for each species *j*.

### Fitting procedure

To fit a protein-DNA interaction model to the binding data, we perform 80 separate regressions and select the set of binding parameters (the “motif”) for which the linear correlation *r* between the predicted number of particles 〈*N*〉 (Eq. 5) for each DNA probe and the target data (MITOMI or PBM fluorescence ratios) is the highest. Each regression starts with a randomly generated motif. At each subsequent step, the motif from the previous step is modified by adding a random value Δ*ε*, drawn from a normal distribution of unit variance, to a single randomly chosen parameter.

When one of the parameters (except *ε*^0^) is modified, some of the other parameters have to be changed as well in order to satisfy the constraints. For example, in the mononucleotide model, if 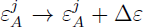, 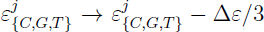 is used in order to offset the shift. In the dinucleotide model, mononucleotide parameters are treated as above, and dinucleotide parameters are modified as follows: if 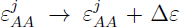, 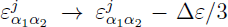 for all dinucleotides that have *A* at either first or second position, and 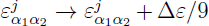 for all other dinucleotides. In general, for words of length *n* we have:

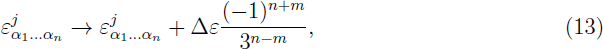

where *m* is the number of letters in common between *α*_1_ … *α*_*n*_ and the word whose energy was shifted by Δ*ε* (*j* is omitted for position-independent parameters).

After modifying the fitting parameters, the new *r* is evaluated. If it is better than the previous one, the new motif is accepted, and the algorithm proceeds to the next regression step. If the new *r* is not an improvement, another trial step is made. If no improvement is found after 200 trial steps, the modification that decreased the correlation least is accepted, and the algorithm proceeds to the next regression step [20]. We call this trial procedure Try2Step; see SI Materials and Methods for the pseudocode. After *T* = 400 regression steps have been made, *r* of the current step is compared to *r* obtained *T* steps ago. If this difference is less than 0.003, the current regression ends and its best *r* and the corresponding final motif are recorded. The motif that achieved the best linear correlation *r*_best_ among 80 separate regression runs is reported as the final prediction.

### Motif Correlation

We find correlations between motifs produced by energy models of arbitrary complexity. First, we convert each energy parameter to probabilities:

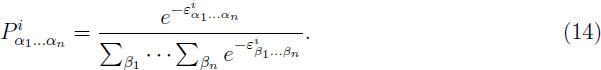

Here, 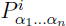 is the probability of the word *α*_1_ … *α*_*n*_ at position *i* (*i* is omitted for position-independent parameters). The information content *R*^*i*^ is then given by the difference between the maximum possible and the observed entropy [21]:

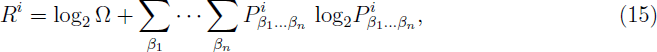

where Ω is the alphabet size (e.g., 4 for mono- and 16 for dinucleotides). Finally, the “height” of each word is defined as

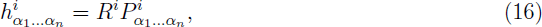

similarly to letter heights in standard PWM logos [21].

We compare two motifs with the same number of energy parameters using word heights. We allow one motif to be offset from the other by up to 4 bases in either direction. We also create a reverse complement of the other motif, testing 18 alignments in total. For each aligment, a linear correlation between the two sets of word heights is computed; the best alignment is that with the maximum correlation. Note that when the two motifs are offset from one another, only the subset of word heights that corresponds to the aligned part is taken into account. Figure S1 shows a histogram of motif correlations among randomly generated mononucleotide models with *L* = 10. To generate energy parameters for random motifs, we start with all parameters set to zero. Then for each letter *α* at each position *i* we draw a number from a normal distribution with zero mean and standard deviation 2. We add this number to the current 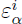 and enforce the constraints using Eq. 13. In the end, we compute word heights via Eq. 16. Using this histogram, we approximate the *p*-value for a mononucleotide motif correlation of *r* as the fraction of correlations ≥ *r*. The *p*-values for dinucleotide motifs are obtained similarly.

### IntervalLogo

We have devised a novel way of displaying parameter confidence intervals in sequence logos, called IntervalLogo. We define the confidence interval for each energy parameter as the range of values of that parameter within which the correlation *r* ≥ 0.98*r*_best_ when the parameter is changed and all the other parameters are held fixed, except to satisfy the constraints. We will demonstrate our procedure using the mononucleotide model; more complex models are treated in a similar manner. Let us say that 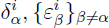 are the values of the energy parameters at the confidence boundary. Here, 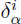 is the parameter modified explicitly, and 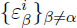 are the other parameters offset to satisfy the constraint (Eq. 13). Given these energies, the probability of each letter at both confidence boundaries is defined by Eq. 14, the information content by Eq. 15, and the word heights 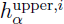 and 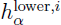 by Eq. 16. By construction, we enforce 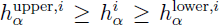, where 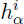 is given by Eq. 16 with best-fit parameters. Note that due to non-linear relationship between energy parameters and word heights it is possible to have 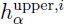 and 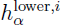 associated with either end of the confidence interval for the energy parameter in question. Occasionally, 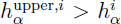 and 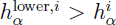, where 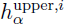 is now associated with the high-energy boundary for the energy parameter, and 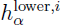 is associated with the low-energy boundary. In this case, we simply set 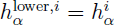. Likewise, if 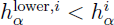 and 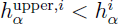, we set 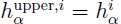.

In the IntervalLogo, the height of each letter is given by 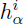; the letters are sorted by their heights 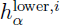 at the *lower* confidence boundary, so that letters with smaller 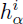 and small errors occasionally appear on top of the letters with larger 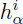 but also larger uncertainties. To show confidence boundaries in the logo, we draw the letter with light, medium, and dark colors (Fig. 1; the choice of the overall color is arbitrary). The light color, representing the larger word height, is painted from the top of the letter down a distance 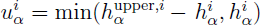 (so that if 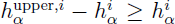, the light color is painted over the whole letter). The dark color, representing the smaller word height, is painted from the bottom of the letter up a distance 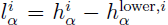. If there is no overlap, the medium color appears between the light and dark regions (Fig. 1A). Otherwise, the light and dark colors are striped in the region of overlap, and the medium color is not visible (Fig. 1B).

**Figure 1:**
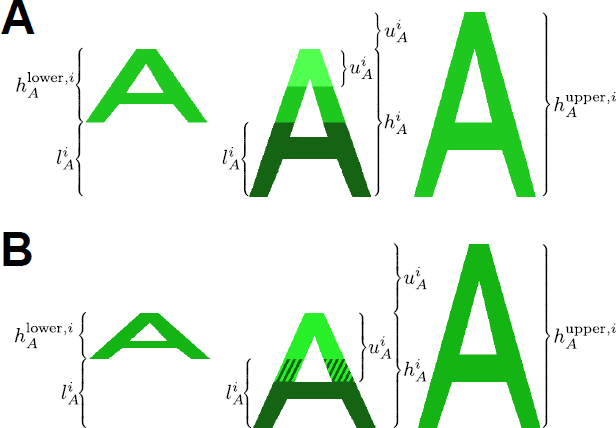
Illustration of color assignments in IntervalLogos. All symbols are defined in Materials and Methods.

### Mononucleotide logos for higher-order models

We can represent higher-order motifs using a conventional mononucleotide sequence logo. To accomplish that, we predict the probability 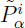 of finding base *α* at position *i* within the binding site, for all possible sites of length *L*. For mononucleotide motifs, 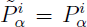 from Eq. 14; for higher-order motifs a recursive calculation is carried out.

Specifically, we need to compute

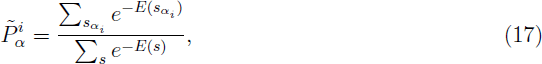

where *s* is a set of all sequences of length *L*, and 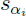 is the subset of these sequences with the nucleotide *α* at position *i*. In order to compute 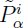 recursively, we first define 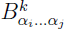, the product of all Boltzmann weights due to substrings of the word *α*_*i*_ … *α*_*j*_ covering position *k*, where *i* ≤ *k* ≤ *j*. 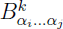 is given by

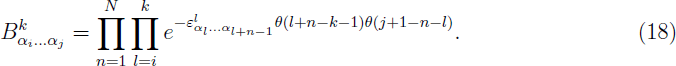

We now introduce the forward partition function

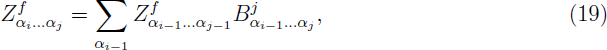

where *j* − *i* = *N* − 2, and the initial condition is that 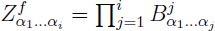 for *i* < *N*. Note that with *k* = *j* and *j* − *i* = *N* − 2, Eq. 18 reduces to 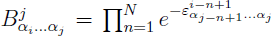. The backward partition function is given by

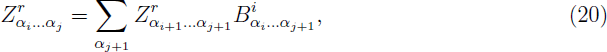

where *j* − *i* = *N* − 2, 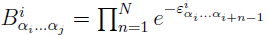, and the initial condition is that 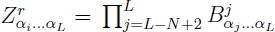 for *i* > *L* − *N* + 1.

We then compute the full partition function from *Z*^*f*^ or *Z*^*r*^:

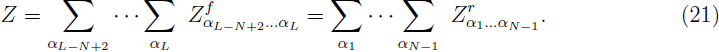

Finally, the marginalized probability is

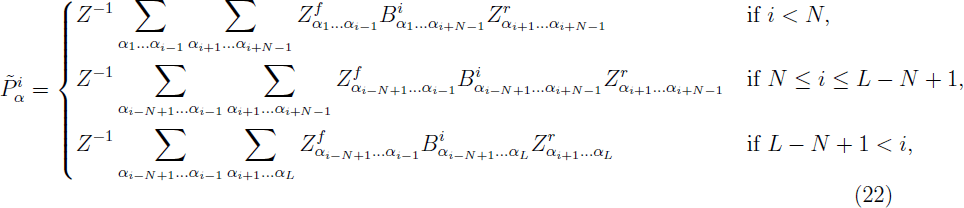

where the 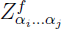 and 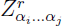 terms are set to 1 whenever *i* > *j*. Specifically, 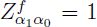 if *i* = 1 and 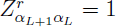 if *i* = *L* in Eq. 22.

## Results

### Overview of the modeling procedure

We have developed an algorithm, BindSter, which predicts TF-DNA binding energetics using a biophysical approach (Fig. 2). The algorithm assigns a binding energy *E*_*i*_ to every site *i* … *i* + *L* − 1 of length *L* on both DNA strands. The energy is computed using a hierarchy of models of increasing complexity (Materials and Methods). The most basic of these, the mononucleotide model, assumes that the contribution of each DNA bp to the total binding energy is independent of the other bps (Eq. 7).

The mononucleotide model can be extended by either including energies of dinucleotides (adjacent base pairs; Eq. 8), or k-mers of maximum length *N* that can occur anywhere within the binding site (Eq. 9). The dinucleotide model is designed to capture sequence specificity of DNA bending, which plays a major role in protein-DNA recognition [22–24] and which is typically described at the level of base-stacking energies [25]. The k-mer model is capable of accounting for sequence-specific experimental biases [19], as well as capturing true sequence preferences for longer motifs. We have also studied a position-independent (PI) model in which protein-DNA energetics is described purely through k-mer contributions (Eq. 11).

We assume thermodynamic equilibrium between proteins and DNA probes. Under this assumption, we compute the average total number of proteins bound to each DNA probe using efficient transfer matrix-like algorithms (Materials and Methods) [13, 14]. Unlike previous approaches [15, 17] which assume low protein concentrations, our algorithm sums over all configurations of DNA-bound proteins allowed by steric exclusion. Furthermore, several species of DNA-binding factors can compete for positions on the DNA probe; this faculty allows us to fit two models of protein-DNA interactions simultaneously in order to detect alternative binding modes.

BindSter regressions start from a set of random points in the parameter space, so that the fits are not biased by initial conditions. Input data comes in the form of the ratio of Cy5 to BODIPY fluorescense intensity for MITOMI [10], and Alexa488 to Cy3 for PBM [12, 26, 27] (Fig. 2B). Each regression attempts to maximize a linear correlation coefficient *r* between measured fluorescense ratios and predicted protein occupancy (Fig. 2C,D). Parameter optimization is carried out using a simple derivative-free algorithm which we call Try2Step [20] (Materials and Methods). Briefly, the algorithm tries out modifications of the current parameter set and accepts the first one that increases *r*; if no such modifications are found after 200 steps, the parameter set that decreased *r* the least is accepted. This offers a possibility of surmounting barriers on the landscape defined by the dependence of *r* on model parameters, although most accepted steps do lead to improvements (Fig. 2C). The regression is stopped once improvements to *r* become too small to matter. After 80 independent regressions, the model which yields the highest *r* is retained as the final prediction (Fig. 2D). Each prediction consists of a set of final model parameters, typically shown as a sequence logo, and of the protein occupancy profile for each DNA probe.

**Figure 2:**
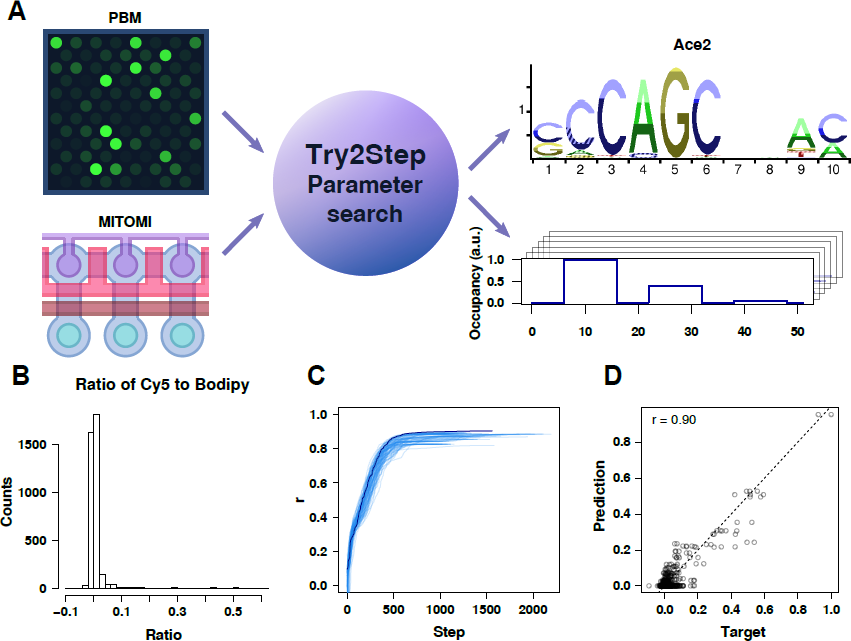
Overview of BindSter, a biophysical algorithm for predicting protein-DNA binding specificity from high-throughput data. (A) MITOMI or PBM data is used as input to a series of Try2Step regressions employed to optimize model parameters. The fitted model (here, the Ace2 mononucleotide model) can be displayed as a sequence logo (e.g., an IntervalLogo; see Materials and Methods for details) and used to compute a TF occupancy profile for each DNA probe. (B) Histogram of the Cy5 to BODIPY ratio, which quantifies the level of TF binding, from the Ace2 MITOMI assay [10]. (C) Multiple regressions on the Ace2 MITOMI data. Each regression attempts to maximize the linear correlation coefficient between probe fluorescense ratios and the average number of particles bound to each probe (see Materials and Methods). The regression leading to the highest correlation is plotted in dark blue. (D) Predicted average number of particles vs. observed fluorescence ratios for each probe from the Ace2 MITOMI assay. The predictions are made using the mononucleotide model.

### Sensitivity of prediction quality to model parameter values

Each BindSter run yields a set of model parameters that fit the data best. However, focusing only on best-fit models overlooks the fact that fit quality may be insensitive to the exact values of some of the fitting parameters. If the dependence of *r* on a given parameter is weak, different algorithms may yield apparently different motifs that would in fact have equivalent predictive power. To address this issue, we have developed a graphical representation of parameter uncertainties, called an IntervalLogo (see Materials and Methods for details). In IntervalLogos, letter heights correspond to predictions based on the best-fit model, as in regular logos. In addition however, color intensities and patterns are used to display the confidence interval associated with each letter (Fig. 1). The intervals are based on varying model parameters one at a time, until *r* drops to 0.98 of its optimal value found in the best-model fit.

### The mononucleotide model and comparison with previous work

We have fit the mononucleotide model to 28 *S. cerevisiae* TFs with MITOMI data [10] (Fig. 3A, Table S1). Our results can be compared with previous fits of a PWM-like model to the same data, which employed MatrixREDUCE [10]. When the same length of the binding motif is used (*L* = 8), our results are very similar to MatrixREDUCE predictions (p=0.31, red circles in Fig. 3A). However, we find that prediction quality improves when motifs are longer (compare crosses (*L* = 10) and circles (*L* = 8) in Fig. 3A), and therefore use *L* = 10 in all subsequent fits. Despite comparable overall predictive power, MatrixREDUCE and BindSter sometimes yield significantly different motifs (Table S1, Fig. 3C), indicating that (a) parameter uncertainty may be significant at some positions; (b) more than one model of binding specificity may explain the same data equally well.

Note that in all MITOMI fits we have used only the 52-bp variable region from each DNA probe; refitting the *L* = 10 mononucleotide model with the entire 70-bp probe sequence results in a modest improvement compared to the result shown in Fig. 3A (violet crosses): the average change in correlation coefficients *μ* is 0.013, with *p* = 1.2*e* − 4. The average motif correlation is 0.86, with motifs produced by poorly-fitting models being more variable.

**Figure 3:**
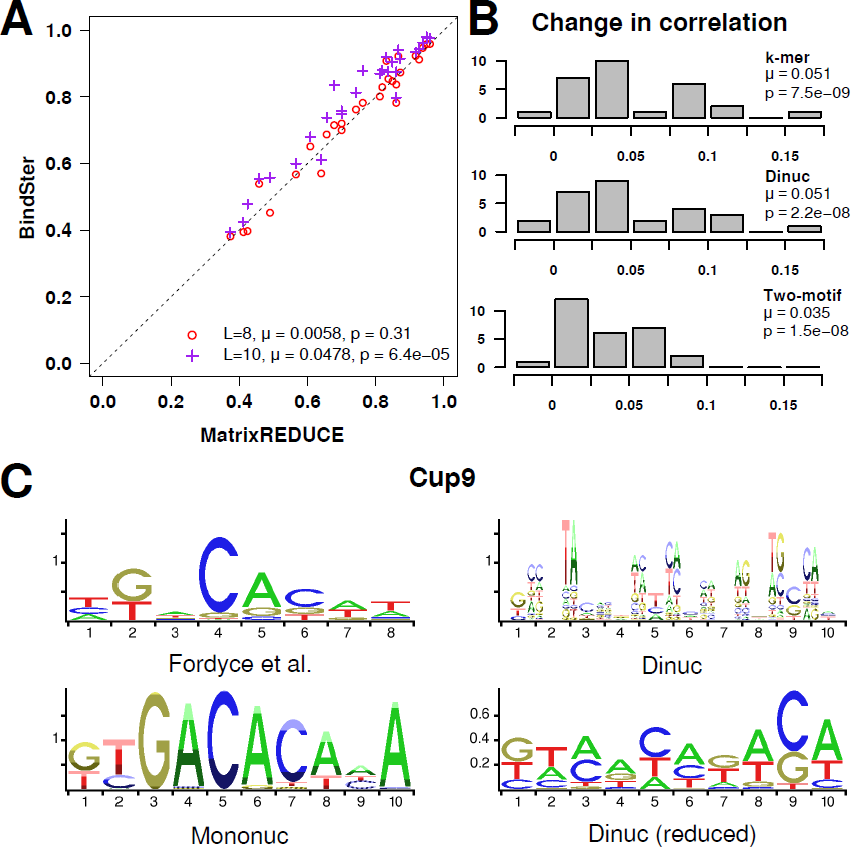
Protein-DNA interaction motifs obtained from MITOMI data. (A) Comparison of MatrixREDUCE [15] linear correlation coefficients [10] with those from BindSter. BindSter fits were done using a mononucleotide model with *L* = 8 (as in MatrixREDUCE) or *L* = 10. The difference between BindSter and MatrixREDUCE linear correlation coefficients averaged over all TFs is denoted by *μ*; *p*-values for the two distributions of correlation coefficients are computed using the Wilcoxon signed-rank test. (B) Histogram of changes in correlation coefficients between extended (k-mer, dinucleotide, and two-motif) and mononucleotide models. The difference between extended and mononucleotide model correlation coefficients averaged over all TFs is denoted by *μ*; *p*-values are computed as in (A). (C) Motif predictions for Cup9. Counter-clockwise from top left: Fordyce et al. mononucleotide logo, BindSter mononucleotide IntervalLogo, BindSter dinucleotide IntervalLogo, the projected logo based on the dinucleotide model (Materials and Methods). See Table S1 for all other TFs and the k-mer model.

### Steric exclusion

One advantage of BindSter is its rigorous treatment of steric exclusion, which makes our algorithm applicable to arbitrary protein concentrations (at low concentrations, multiple proteins bound to the same DNA probe are rare and steric exclusion is not important). The additional computational effort is modest since we use an efficient recursive algorithm to calculate DNA probe occupancies (Materials and Methods). In order to investigate the extent to which neglecting steric exclusion affects the quality of predictions, we have compared the mononucleotide model to a simple model without steric exclusion, which treats each binding position as independent of all the other positions on the DNA probe (Materials and Methods). We find that rigorous treatment of steric exclusion has minimal effect on the predictive power of the model (Fig. S2). Both the correlation coefficients (Fig. S2A) and the motifs (Fig. S2B) are very similar, although there are occasionally minor differences in probe occupancy profiles. For example, Rox1 profiles are somewhat different with and without steric exclusion (Fig. S2C), probably due to the preference for T/C at position 2 in the latter fit (Fig. S2B). It appears that in MITOMI experiments the protein concentrations are sufficiently low, making steric exclusion a minor factor.

### Higher-order models of protein-DNA energetics

The mononucleotide model may not fully capture the complexity of amino acid – DNA base pair interactions at the protein binding interface [28], including the role of DNA shape in protein recognition [24]. To determine whether energetic contributions of longer motifs improve predictive performance, we have tested several extensions of the basic mononucleotide model (Fig. 3B, Fig. S3, Table S1, Table S2; see Materials and Methods for model definitions). The extended models yielding the best improvements are the *N* = 3 k-mer model (where *N* is the maximum word length) and the dinucleotide model (top two panels in Fig. 3B; Table S1). Interestingly, the *N* = 2 k-mer model has significantly less predictive power (Fig. S3, top panel).

Either *N* = 2 or *N* = 3 position-independent model fit simultaneously with the mononucleotide one does not substantially improve performance (Fig. S3, two middle panels). Thus there is scant evidence in MITOMI data for nearly non-specific binding driven by preferences for short sequence motifs, regardless of their position within the binding site. Finally, whereas fitting two mononucleotide models simultaneously is beneficial (two-motif model in Fig. 3B, bottom panel; Table S2), fitting one mononucleotide parameter set first and then fitting the other one on the residual signal is not (secondary motif model in Fig. S3, bottom panel; Table S2). Since the parameter spaces for these two models are equivalent, the issue lies with convergence: simultaneous fits achieve better optimization than sequential ones, probably because of greater flexibility afforded by a higher-dimensional optimization in the former case. Correspondingly, secondary motifs are often less specific and have little predictive power by themselves (cf. *r*_alone_ values for the secondary model motif_2_ in Table S2). In contrast, motif_2_ fits in the two-motif model are more predictive and the logos are often related to motif_1_ and mononucleotide model fits (Table S2). The differences between motif_1_ and motif_2_ predictions may sometimes be interpreted as alternative binding modes. For example, in the case of Gcn4 motif_1_ is close to the symmetric AP-1 site sequence (TGA(C/G)TCA) which corresponds to homodimeric binding [29]; motif_2_, on the other hand, may reflect monomeric binding to the TCA half-site (note however that C at position 7 has a large uncertainty associated with it).

Interestingly, motif specificity (as seen in the heights of logo letters) does not necessarily confer predictive power [19]. For example, the mononucleotide model prediction for the TF Cup9 is significantly more specific than the MatrixREDUCE one (compare two left panels in Fig. 3C), but their linear correlation coefficients are fairly similar (*r* = 0.56 and *r* = 0.49, respectively). The dinucleotide model, which provides a substantial improvement (*r* = 0.73), yields a motif that appears less specific still, although there are several prominent dinucleotide contributions (Fig. 3C, two right panels). Note that the the dinucleotide model’s projected logo in Fig. 3C is constructed using the full dinucleotide parameter set, as described in Materials and Methods.

In general, it is challenging to associate improvements in performance of higher-order models with a particular physical mechanism. For example, in the case of Cup9 the k-mer and dinucleotide models yield very similar correlation coefficients (0.71 and 0.73, respectively; Table S1), yet their physical underpinnings are completely different. The dinucleotide model is designed to capture DNA bp stacking energies, whereas the k-mer model assigns energies to short words with lengths of 2 and 3 nucleotides independently of their position within the binding site, reflecting global preferences for these short motifs. Nonetheless, in the scatter plot of predicted vs. observed DNA probe occupancies (Fig. S4A), predictions of both models co-localize for high-occupancy probes, and are more accurate compared with the mononucleotide model. Furthermore, predictions of both higher-order models correlate equally well with the mononucleotide model and again co-localize for high-occupancy probes (Fig. S4B). A direct comparison of k-mer and dinucleotide models reveals that they predict very similar distributions of probe occupancies (*r* = 0.90; Fig. S4C). Thus the same signal is captured using two distinct higher-order models, which extend the mononucleotide model in different ways.

The above observations lead us to conclude that modifying basic mononucleotide energetics is more beneficial than introducing additional binding species. Indeed, according to our fits there is relatively little evidence that TFs switch between binding modes and in addition to the primary bind an alternative motif which could be characterized by either word counts (position-independent models) or another mononucleotide parameter set (two-motif and secondary motif models). The most successful approach of this type, the two-motif fit, typically yields primary and secondary motifs that are clearly related (Table S2).

### Cross-validation of BindSter fits

For every TF, the MITOMI dataset provides approximately 3700 DNA probe occupancy measurements; the full dataset is divided into two complete replicates and an incomplete replicate which contains a subset of probes from the complete one [10]. Thus the number of free parameters in all our models (Materials and Methods) is significantly less than the total number of available measurements. Nonetheless, we have cross-validated our mononucleotide fits by comparing the models trained on first and second complete replicates with each other, and with the model trained on all available data (Fig. S5, Table S3, Table S4). Models trained on single replicates produce distributions of probe occupancies that are highly correlated with the full model (Fig. S5A) and have similar predictive power against the data (Table S3). The fitted model parameters also tend to be similar, as measured by motif correlation coefficients against the mononucleotide model trained on the entire dataset (Fig. S5B, Table S4). As a rule, predictions with high correlation coefficients against the data tend to yield nearly identical motifs (see e.g. Rox1 and Sko1 examples in Fig. S5C); motifs vary more for lower-quality predictions (see e.g. the Cad1 example in Fig. S5C).

It is also possible that higher-order models fit the data better simply because they have more fitting parameters. However, there is no clear connection between the number of additional parameters and model performance. For example, the best-performing models in Fig. 3B and Fig. S3 are the dinucleotide model (112 independent parameters compared with 31 for the mononucleotide model) and the k-mer model (67 independent parameters with *N* = 3). Despite many more parameters in the dinucleotide model, its performance is quite similar to that of the k-mer model, as discussed above.

The comparison between the k-mer and mononucleotide + PI models is particularly instructive. These two models are parameterized nearly identically, but k-mer models greatly outperform their PI couterparts. The physical mechanisms of protein-DNA interactions described by the two models are very different. In the mononucleotide + PI model, TFs bind in two distinct modes (Materials and Methods). In the k-mer model, there is a single binding mode whose sequence preferences are described by a combination of position-specific mononucleotide parameters and position-independent contributions from longer words. Thus merely adding parameters will not significantly improve predictive performance unless the underlying physical model captures essential aspects of protein-DNA energetics.

Since the dinucleotide model has the most fitting parameters, we have also cross-validated it directly (Table S3, Table S4). As with the mononucleotide model, predictions of DNA probe occupancies are largely consistent across the two models trained on separate replicates, with slight loss of predictive power which becomes more pronounced if the correlation coefficients from all models (replicate 1, replicate 2, and full) are low overall (Table S3). However, dinucleotide motifs fitted on replicates are much more variable than their mononucleotide counterparts (Table S4). For example, Ace2 models fit on separate replicates have the same predictive power as the full model. Nonetheless, the corresponding motifs are only correlated with the full model motif at *r* = 0.51 (*p* < 10^−6^) and *r* = 0.68 (*p* < 10^−6^). For Met31, the dinucleotide motif fit on replicate 1 is correlated with the full motif at *r* = 0.85 (*p* < 10^−6^), whereas the replicate 2 fit yields *r* = 0.37 (*p* = 2.3 × 10^−5^). Interestingly, both motifs predict DNA probe occupancies equally well (*r* = 0.82 and 0.80, respectively), and comparably to the full model prediction (*r* = 0.85). Overall, these observations underscore the fact that multiple models, and especialy multiple higher-order models, can fit MITOMI data equally well.

### Comparison of PBM and MITOMI-based predictions

Predictions of protein-DNA binding specificity should not depend on the experimental platform being used to measure protein-DNA interactions. In order to compare the robustness of BindSter predictions across MITOMI and PBM platforms, we have carried out mononucleotide, dinucleotide, and k-mer fits for 12 TFs for which both PBM and MITOMI measurements are available (Table S5). We have also compared our results with previously published BEEML-PBM PWM-like fits [16, 17].

One essential difference between BindSter MITOMI and PBM fits is the use of location bias in the latter. The location bias in our model is a quadratic energy contribution with two free parameters per DNA-binding species (Eq. 12). It is designed to make binding sites closer to the glass slide less favorable, due to their lower accessibility for TFs [19]. Indeed, although the coefficients of the quadratic function are not constrained in any way in our fits, for most TFs they fit to values that impose energetic penalties on the proximity of the protein to the glass slide (Fig. S6A). Including location bias into PBM fits leads to sizable improvements in prediction quality for mononucleotide and k-mer models (Fig. S6B). In contrast, including location bias into MITOMI fits leads to *μ* = 0.013, several times smaller than the improvements observed with PBM data. Although including location bias increases predictive power of PBM fits, mononucleotide models fit with and without it tend to yield similar motifs (Fig. S6C; average motif correlation for the 5 TFs in the upper panel of Fig. S6B is 0.92).

On average, our mononucleotide predictions outperform those from the BEEML-PBM algorithm (Fig. 4A, Table S5). As with MITOMI data, k-mer and dinucleotide models provide statistically significant improvements compared to mononucleotide fits. Consistency between PBM and MITOMI-based motifs (as measured by motif correlations) improves with overall prediction quality on PBM data; in several cases higher-order models recover motifs that, when projected, are closer to the MITOMI-based mononucleotide motifs than mononucleotide models inferred from PBM data (Fig. 4B). We also note that BEEML-PBM motifs have the highest average correlation with BindSter MITOMI motifs on this dataset. For example, the core GACACA Cup9 motif seen in both BEEML-PBM and MITOMI-based mononucleotide fits is only recovered at the dinucleotide or k-mer level with BindSter PBM fits (Fig. 4C, Table S5). This may be due to substanial k-mer bias present in PBM data [19], which may also be accounted for indirectly in the dinucleotide model.

Predictive power of MITOMI-trained models on PBM data is modest overall, although it does improve if location bias is included (Table S6). As a rule, PBM-based fits yield lower correlation coefficients and partial or less specific motifs. For example, for Bas1 neither BEEML-PBM nor BindSter fits recover the highly predictive motif (*r* = 0.83) inferred from MITOMI data, and even the high-quality prediction of the canonical Cbf1 motif with PBM-based mono- and dinucleotide models has less information content than the corresponding MITOMI result (Fig. 4C, Table S5). Note however that with MITOMI data the symmetric CACGTG motif is replaced with GACGTG, perhaps as a sign of an alternative binding mode; PBM fits on the other hand yield the expected symmetric motif. Interestingly, the Cbf1 motif becomes significantly more specific when the potential k-mer bias is taken into account (Table S5).

**Figure 4:**
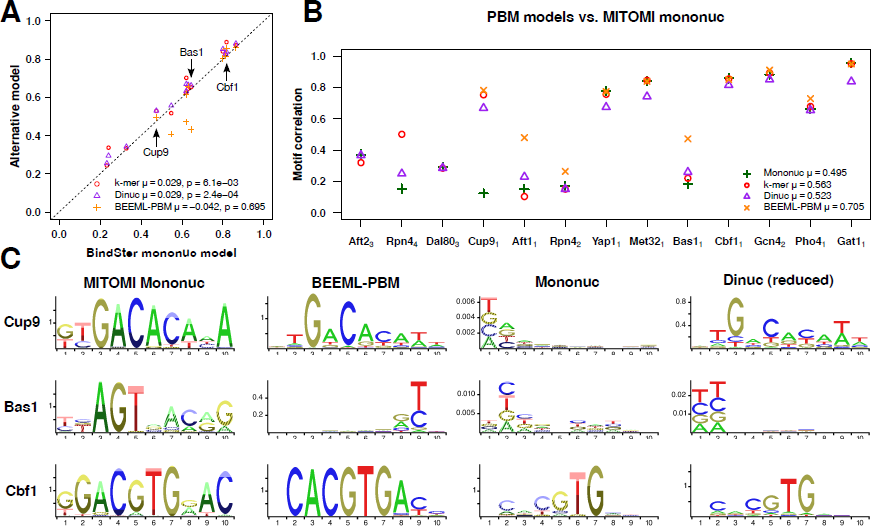
Protein-DNA interaction motifs obtained from PBM data. (A) Comparison of linear correlation coefficients predicted using the mononucleotide model with BEEML-PBM [16, 17], dinucleotide, and k-mer models. All models are fit on PBM data. The average difference between linear correlation coefficients in each comparison is denoted by *μ*; *p*-values for the two distributions of correlation coefficients are computed using the Wilcoxon signed-rank test. (B) Motif correlations between the mononucleotide BindSter model fit on MITOMI data and three BindSter models (mononucleotide, k-mer, and dinucleotide) as well as the BEEML-PBM model fit on PBM data. Dinucleotide and k-mer motifs are projected onto the mononucleotide parameter set for this comparison (see Materials and Methods). All motif correlations are sorted by the linear correlation coefficient obtained by fitting the mononucleotide BindSter model to PBM data; *μ* indicates the average correlation coefficient for each model. Subscripts on TF names denote the source of the PBM data (see Table S5 for details). (C) Representative logos for MITOMI mononucleotide, BEEML-PBM, PBM mononucleotide, and PBM dinucleotide fits (see Table S5 for complete results). The dinucleotide model is represented by its projected logo.

## Discussion

We have developed a biophysical algorithm, BindSter, for predicting protein-DNA binding energetics from high-throughput *in vitro* measurements of total protein occupancies on a set of DNA probes. We have focused on MITOMI, a technique that uses a microfluidic device to measure binding interactions at thermodynamic equilibrium [9, 10]. The MITOMI binding data is thus well-suited for computational modeling based on equilibrium statistical mechanics, although the MITOMI approach is less high-throughput than PBMs [12, 26, 27, 30].

Our computational framework differs from previously published biophysical models of protein-DNA interactions [15–17] in two important aspects: (a) It employs a rigorous procedure in which all sterically allowed configurations of TFs bound to DNA probes are taken into account; (b) It is designed to accommodate a variety of protein-DNA interaction energy models. Thus, in contrast to previous work, BindSter is able to provide a quantitative treatment of overlapping binding sites and multiple DNA-binding species (as may occur if the TF switches between dimeric and monomeric binding, or employs distinct binding interfaces to interact with different classes of sites). The built-in flexibility with respect to describing protein-DNA energetics has allowed us to explore a hierarchy of models of increasing complexity.

The most basic mononucleotide model, in which each nucleotide in the binding site contributes independently to the total protein-DNA binding energy, has predictive power on MITOMI data comparable to that of MatrixREDUCE [10]. However, many BindSter motifs look substantially different (Table S1). Some of this discrepancy may be attributed to the fact that the models are insensitive to the values of a subset of fitting parameters. Such parameters could therefore change within a sizable range with little consequences for the quality of the fit. To account for this possibility, we have developed a novel way of graphically displaying the uncertainty of model parameters, called IntervalLogos.

IntervalLogos are constructed as regular logos, but with color intensities and patterns of the letters used to indicate the confidence interval associated with each parameter as it is varied independently (Materials and Methods; Fig. 1). However, visual inspection of mononucleotide and MatrixREDUCE motifs in Table S1 shows that even after accounting for parameter uncertainty the motifs remain related but distinct in many cases. Thus there is degeneracy among motifs that can explain MITOMI data equally well; moreover, there is no clear correlation between the specificity of a given motif and its predictive power. Finally, the role of steric exclusion is negligible in MITOMI experiments, probably due to low protein concentrations which make multiply bound proteins and competition for the same DNA sequence unlikely (Fig. S2).

Among seven extensions of the mononucleotide model that we have tested, the dinucleotide and k-mer models result in most significant improvements to prediction quality (Fig. 3B, Fig. S3). Both of these models extend the mononucleotide description of proteinDNA energetics by including contributions of longer words. In contrast, models that test for the presence of alternative binding motifs fair less well, indicating that TFs in our dataset tend to have a single predominant binding mode. The most successful model of this type, in which two mononucleotide motifs are fit simultaneously to the data, does result in a reasonable improvement (Fig. 3B); some of the fitted model pairs can be interpreted as evidence of dimeric vs. monomeric binding, or affinity for different classes of sequence motifs (Table S2). However, in general it is challenging to associate an observed improvement in model performance with a particular physical mechanism of protein-DNA interactions. For example, k-mer and dinucleotide models tend to produce correlated predictions for high-occupancy DNA probes despite the fact that they are supposed to describe distinct aspects of protein-DNA energetics and/or experimental biases (Fig. S4).

Finally, we find that fits on PBM data typically yield substantially different motifs, depsite the fact that their predictive quality is as good or better than previously published BEEML-PBM results [16, 17] (Fig. 4, Table S5). Motif correlations between MITOMI and PBM-fit models are modest overall, although they do improve with the quality of the PBM fit (Fig. 4B). Correspondingly, the ability of MITOMI-trained data to predict the distribution of probe occupancies (fluorescent intensities) observed in PBM experiments also tend to be modest (Table S6). One potential explanation is limited applicability of thermodynamic models to the analysis of PBMs. Another possibility is experimental biases other than the k-mer bias and the location bias that were explicitly taken into account in BindSter modeling.

